# Assembly Arena: Benchmarking RNA isoform reconstruction algorithms for nanopore sequencing

**DOI:** 10.1101/2024.03.21.586080

**Authors:** Mélanie Sagniez, Anshul Budhraja, Bastien Paré, Shawn M. Simpson, Clément Vinet-Ouellette, Marieke Rozendaal, Martin A. Smith

## Abstract

Resolving the transcriptomes of higher eukaryotes is more tangible with the advent of long read sequencing, which greatly facilitates the identification of new transcripts and their splicing isoforms. However, the computational analysis of long read RNA sequencing data remains challenging as it is difficult to disentangle technical artifacts from *bona fide* biological information. To address this, we evaluated the performance of multiple leading transcriptome assembly algorithms on their ability to accurately reconstruct RNA transcript isoforms. We specifically focused on deep nanopore sequencing of synthetic RNA spike-in controls (Sequins™ and SIRVs) across different chemistries, including cDNA and direct RNA protocols. Our systematic comparative benchmarking exposes the strengths and limitations of the different surveyed strategies. We also highlight conceptual and technical challenges with the annotation of transcriptomes and the formalization of assembly quality metrics. Our results complement similar recent endeavors, helping forge a path towards a gold standard analytical pipeline for long read transcriptome assembly.

## Introduction

Long read sequencing greatly improves transcriptome profiling via its ability to qualify full-length RNAs, helping resolve complete exon chains and unannotated biological features, such as long non-coding RNAs, which often contain repetitive sequences (Byrne et al. 2019; Lagarde et al. 2017). These technologies present a distinct advantage over short-read RNA sequencing (RNA-seq) that require a well-annotated reference and struggle to resolve long-range splicing dependencies. The utilization of long reads holds promise for expanding the repertoire of known RNA transcripts, improving the quantification genes and splicing at the isoform level while contributing to a more detailed portrait of the transcriptome.

Recent studies have compared software for estimating transcript abundances in long read sequencing data (Dong et al. 2023; Pardo-Palacios et al. 2023b) but few reports have detailed the effectiveness of tools for RNA isoform discovery and qualification using long reads. Transcriptome assembly is an economical method for characterizing gene expression of model (Yang et al. 2021; Kuo et al. 2020) or non-model organisms (Kang et al. 2021; Zagorščak and Petek 2021; Matra et al. 2023) and for identifying novel RNAs in diseases such as cancer (Fang et al. 2021; Verma et al. 2015).

While the definition of de novo assembly encompasses many meanings, we categorize the existing assemblers into 3 distinct overall strategies: (i) guided strategies, which combine sequencing data aligned to a reference genome and an existing reference gene annotation; (ii) *de novo* strategies thah rely solely on reads aligned to a reference genome to reconstruct a transcriptome; and (iii) *ab initio* strategies, which require only sequencing reads (often used for non-model species without a genome assembly). We compared 12 tools that assemble transcripts from long reads according to one (or more) of the above mentioned categories. Some tools (i.e. Bambu, Flair, isoQuant, and Stringtie2) fit into two strategies–namely ‘guided’ and ‘*de novo*’–as they allow the optimal input of a reference transcriptome annotation, such as RefSeq (O’Leary et al. 2016) or Gencode (Frankish et al. 2019), typically in gene transfer file (.gtf) file format.

Bambu (Chen et al. 2023) is a “context aware quantification tool” for long reads. It is an R package that works by using genome-aligned reads (in .bam format) to quantify known and putative novel genes. Bambu is intended to be used as a transcript quantification tool and not a transcriptome assembler. However, it has been included herein as it has been reported in recent studies considering novel transcript identification (Dong et al. 2023; Pardo-Palacios et al. 2023b). The Bambu pipeline has a feature that allows for *de novo* transcript discovery by estimating a novel discovery rate (NDR) which optimizes the false discovery rate in a reproducible manner across samples and analyses. The algorithm annotates reads using assigned read-classes to label splice junctions as full or partial overlaps, thereby producing an intermediate annotation output for the transcriptome. When provided with a transcriptomic annotation (the above-mentioned ‘guided mode’), Bambu outputs all isoforms of that given annotation while adding discovered transcripts below the NDR limit.

Full-Length Alternative Isoform analysis of RNA (or FLAIR) (Tang et al. 2020) is a python workflow created for differential isoform expression analysis that also includes a transcriptome assembly module. It can integrate matched short-reads alongside long-reads to correct errors in the latter and improve accuracy. FLAIR organizes transcripts into groups based on splicing patterns, then further groups these reads by analyzing transcription start sites (TSSs) and end sites (TESs) separately. This is followed by collapsing the grouped reads into the most common isoform within each window for TSS and TES, respectively. FLAIR then aligns these reads to the initial isoform groups. Each putative RNA isoform is then considered to be a valid and observed transcript depending on its observed coverage. FLAIR is composed of several modules that can be run consecutively or independently; thus enabling both *de novo* and *guided* assembly strategies.

Full-Length Analysis of Mutations and Splicing (FLAMES) is a long-read transcriptome analysis tool developed for single-cell analyses that also includes a pipeline for bulk RNA-seq analysis (Tian et al. 2021). FLAMES initially groups reads in a similar manner to FLAIR. However, it introduces a unique approach to modeling the probability of read truncation by incorporating a linear model of isoform length, which leverages the concept that longer isoforms are more likely to have truncated reads with incomplete 5′/3′ ends (especially in the case of single cell RNA-seq droplet-capture protocols where low input material and multiple rounds of PCR are employed). Following this, the sequence of each polished transcript serves as an updated reference for direct realignment of input reads using Minimap2 (Li 2021). During this realignment, transcripts with insufficient coverage are discarded. Reads are then assigned to transcripts based on alignment scores, fractions of reads aligned, and transcript coverage, resulting in an output of isoform-level counts.

IsoQuant (Prjibelski et al. 2023) is a long read transcript annotation tool, written in python, that uses intron graphs to reconstruct transcripts with and without a reference annotation. In the graph, two vertices of splice junctions are connected by a directed edge––if two splice sites are found one after the other in a read, they will be connected in the graph. In the case where a reference annotation is provided, reads are assigned to known isoforms by approximate intron-chain matching to account for error-prone reads. Like Bambu, when provided with a reference annotation, IsoQuant outputs all transcripts from the reference in addition to new transcripts, as well as another assembly exclusively containing isoforms expressed in the input data.

Mandalorian (Volden et al. 2023) is a long-read transcript identification tool also tested in the LRGASP benchmarking project (Pardo-Palacios et al. 2023b). It takes as input accurate full-length transcriptome sequencing data (adaptor and poly-A tail trimmed as well as reoriented reads) and is composed of 5 modules, by default, sequentially executed: (i) alignment to genome; (ii) alignment clean-up; (iii) locus by locus grouping and consensus assembly; (iv) alignment to generate isoform models and group by gene; (v) quantification. Mandalorion is reported to be among the strongest methods for transcript discovery but was only tested on PacBio HiFi and Oxford Nanopore R2C2 reads (both of which employ consensus base calling) for increased sequence accuracy (Pardo-Palacios et al. 2023b). Consequently, the authors of Mandalorion have only tested their tool on such data and recommend using Flair for regular nanopore reads.

StringTie (Pertea et al. 2015) is a *de novo* transcriptome assembly and quantification tool written in C/C++ that operates on genome-aligned files from short- or long-read RNA-seq data. The tool implements a network flow algorithm on the assembled contiguous stretches of transcripts called ‘super-reads’. This algorithm determines the path with the maximum coverage, which allows for adjustments to abundance estimations and produces a more fitting transcript abundance result. The most recent update to the tool, StringTie2, has reduced memory requirements and removed a great percentage of false-positive spliced alignments (Kovaka et al. 2019).

Technology Agnostic LOng-read aNalysis (or TALON) (Wyman et al. 2020) is an annotation dependent tool that analyzes long-read transcriptomes across datasets. The recommended first step in the pipeline is to correct the non-canonical junctions using the python tool “TranscriptClean” (Wyman and Mortazavi 2019). This is followed by the python and R based TALON framework, which involves labeling potential internal priming events, annotating transcript models, recording the transcript abundance, filtering transcript models using biological replicates and ultimately generating a custom transcriptome, used to produce the final transcript and gene count tables. An isoform is considered only when near-alike reads are found above a certain threshold count, in multiple replicates. TALON does not use read clustering or a consensus sequence in an attempt to prevent introduction of incorrect input for its transcript model. While capable of processing a single sample, TALON is meant to process technical/biological replicates (Amarasinghe et al. 2020)–a common occurrence in RNA-seq analyses. TALON has been known to produce a few fallacious transcript predictions in simulated long-read data (Kuo et al. 2020).

*Ab initio* tools are not as commonly used as *de novo* or *guided* ones and, therefore, have not been subject to thorough independent benchmarking. RATTLE is a ‘reference-free reconstructor and quantifier of transcriptomes’ as described by the developers of the C++ and python based tool (de la Rubia et al. 2022). It works by clustering raw reads using a two-step greedy approach, which produces read clusters that should belong to the same gene. These ‘gene clusters’ are further classified into ‘transcript clusters’ by identifying internal gaps in the read sequences. Error correction and polishing of these clusters produces consensus sequences and matched abundance estimations. Using simulated and spike-in synthetic controls, RATTLE’s reference-free assembly algorithm seems to perform better with dRNA data than cDNA(de la Rubia et al. 2022). Although RATTLE seems to perform on par with reference-based assemblers, it is unable to distinguish different isoforms when exon sizes are below a certain limit (de la Rubia et al. 2022). The authors have favorably compared RATTLE to similar clustering algorithms, namely CARNAC-LR and isONclust.

IsONclust (Sahlin and Medvedev 2020) is a greedy search algorithm, implemented in python, that uses minimizers in order to determine similarity and cluster reads together. This approach follows three simple steps for each read: (i) assigning certain reads as ‘representative reads’ of potential clusters and maintaining a hash table of these reads, queried using minimizers; (ii) assigning reads to the cluster with maximal shared minimizers; (iii) conditionally assigning other reads to existing clusters if the similarity is above a certain threshold, otherwise creating a new cluster with the unclassified read as a new representative. Oxford Nanopore has optimized this program and produced the C++ implementation, isONclust2 (https://github.com/nanoporetech/isONclust2) . IsONclust2 was made in collaboration with the authors of isONclust (Kristoffer Sahlin and Paul Medvedev), which resulted in a tool with improved efficiency that takes strandedness into consideration, as well as additional features.

RNA-Bloom (Nip et al. 2020) is a Java based assembly algorithm originally designed for single-cell short-read transcriptome assembly combining bloom filters with De Bruijn graph assembly. This approach to *ab initio* transcriptome assembly was improved upon and applied to long-read bulk sequencing data with RNA-Bloom2 (Nip et al. 2023). The latter uses digital normalization with strobemers–a strategy reported to be less sensitive to mutations than k-mers–before assembling and polishing unitigs (Sahlin 2021). The authors have compared RNA-Bloom2 to RATTLE and found their tool to be faster, less memory intensive while reaching a higher recall and a lower false discovery rate (Nip et al. 2023).

The diversity of recently developed transcriptome assembly tools and the increasing adoption of long read sequencing technologies reinforce the need to distinguish between analytical artifacts (e.g., erroneous isoform detection) and experimental ones (e.g., fragmented RNA, reverse transcription drop off, internal priming, etc). Here, we describe a comprehensive and independent comparative performance benchmark of the aforementioned transcriptome assembly tools. We assessed algorithmic performance with two tools for comparative isoform qualification, SQANTI3 (Tardaguila et al. 2018) and GFFcompare (Pertea and Pertea 2020). SQANTI3 has recently been employed to evaluate long-read transcriptome tools in the Long-read RNA-Seq Genome Annotation Assessment Project (LR-GASP) study (Pardo-Palacios et al. 2023b). GFFcompare derives from Cuffcompare, a utility program from the well-known RNA-seq analysis suite Cufflinks (Pertea and Pertea 2020). We present a comprehensive overview of the strengths and weaknesses of *de novo* transcriptome assemblers for long reads, with the ultimate objective of helping the scientific community make informed decisions when selecting the most appropriate strategy for their isoform-level transcriptome analyses.

## Results

Transcriptome assembly is a crucial step in deciphering the functional landscape of a cell or organism. We evaluated most available long read transcriptome assembly algorithms encompassing 3 assembly paradigms (*guided*, *de novo* & *ab initio*) using two distinct *in vitro* transcribed RNA spike-in controls, namely Sequins™ (Hardwick et al. 2016) and Spike-in RNA variants (or SIRVs) (Lexogen SIRV Set 4), and four distinct nanopore sequencing chemistries, including the latest direct RNA sequencing protocol (**Table 1**, **Figure 1**).

**Figure 1.**
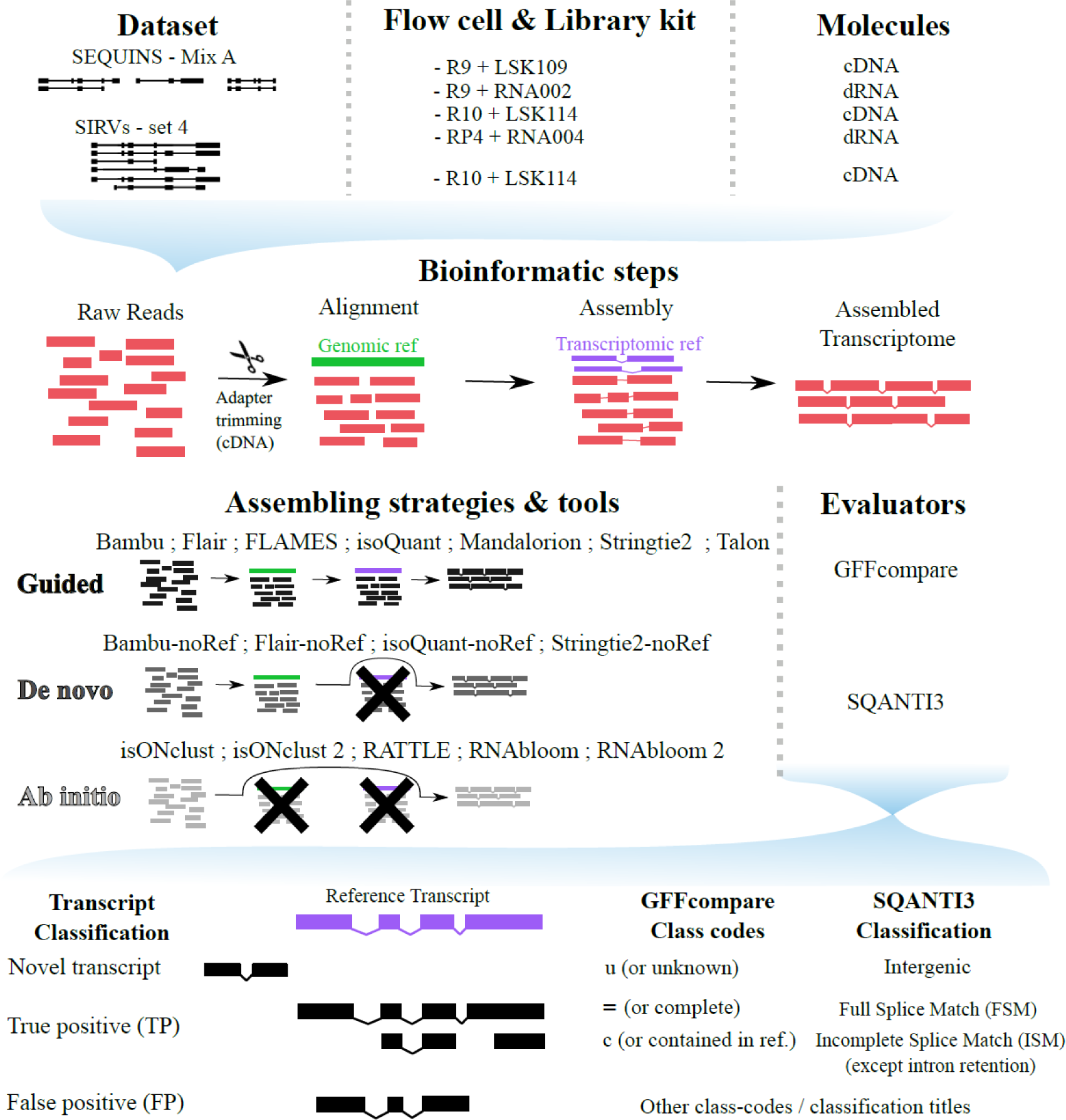
Overview of the study design for cDNA and dRNA assembly generation and quality assessment. LSK: Ligation sequencing kit; cDNA: complementary DNA; dRNA: direct RNA.

**Table 1.** Sequencing data.

| Dataset | Input | Library preparation protocol | Flowcell & ID | Library molarity | Sequencer | Runtime | Basecalling* | Reads (Pass/Fail) | Input number of reads |
| --- | --- | --- | --- | --- | --- | --- | --- | --- | --- |
| LSK109_sequins | 15ng RNA Sequins™ mixA | PCS109 RT protocol + LSK109 | R9.4.1 FAN54376 | 50 fmol | GridION | 72h | Guppy6.0.1_sup | 6,455,950 P+F | 5,2912,936 |
| LSK114_sequins | 15ng RNA Sequins™ mixA | PCS109 RT protocol + LSK114 | R10.4.1 FAV99142 | 20 fmol | PromethION | 12h | Guppy6.4.1_sup | 5,112,902 P+F | 4,262,005 |
| LSK114_SIRVs | 0.0375ng SIRV set-04 + 125ng human RNA | PCS109 RT protocol + LSK114 | R10.4.1 PAK95982 | 18 fmol | PromethION | 72 hours, wash + reload (18 fmol) after 29h | Guppy6.5.7_sup | 42,764** P+F | 42,764 |
| RNA002_sequins | 300ng RNA Sequins™ mixA | RNA002 | R9.4.1 PAM58324 | 20 fmol | PromethION | 89h30m | Guppy6.0.7_hac | 877,710 P+F | 877,710 |
| RNA004_sequins | 4 samples: 15ng RNA Sequins™ mixA 1.5µg human RNA | RNA004_beta-v2 | RP4_beta-testing PAO83093 PAO83456 PAO84072 PAO96683 | 20 fmol | PromethION | 68h, wash + reload (20 fmol) after 44h15m | Guppy6.4.6_sup | 162,644 P | 162,644 |

\*sup = super high accuracy ; hac = high accuracy
\*\*Sequins™ reads retrieved (full-length + rescued), after Pychopper was executed on human + Sequins™/SIRV reads

As qualitative assessment of assemblies is somewhat subjective, we opted to utilize and compare two different tools with distinct yet similar classification categories and scoring metrics, namely GFFcompare and SQANTI3. These were used to compare the assembled isoforms against ‘truth set’ annotations provided by the respective manufacturers of the spike-in controls. N.B. biological samples were purposefully omitted herein given their lack of an absolute truth and the difficulty in distinguishing biological variation from analytical artifacts. Because long-read sequencing sometimes results in low confidence sequences at the 3’ and 5’ ends, we defined a True Positive as a perfect exon-chain match or contained in reference (“c” classcode for GFFcompare and “ISM” for SQANTI3). Precision and sensitivity were calculated for each tool and chemistry combination based on individual classifications of each assembled transcript (see Methods for details). In this context, precision reflects the integrity of the transcriptome, while sensitivity reflects its completeness.

As expected, *guided* strategies perform better than others in sensitivity and/or precision (**Figure 2 & Supplemental File 1**). It is important to note that the most highly performing tool, Bambu, systematically outputs all transcripts from the reference provided to it, in addition to the potential *de novo* transcripts that it assembles, hence surpassing every other tool in both precision and sensitivity in all datasets. Isoquant and Stringtie2 tend to perform comparably, with IsoQuant a step ahead in precision with the Sequins™ controls. Interestingly, the performance of reference-guided IsoQuant appears much inferior on the SIRV controls, which generally contrast less gene diversity but more alternative splicing isoforms than Sequins™. However, IsoQuant consistently performs well across both cDNA and dRNA. Unlike Bambu, which systematically outputs all transcripts irrespective of their expression status, IsoQuant provides the option to only produce transcripts demonstrably expressed within the input dataset (similar to Stringtie2), which was considered herein. Stringtie2 also performs decently in its *guided* mode, although its accuracy decreases notably in dRNA datasets (RNA002 and RNA004 in Figure 2B). For TALON, incorporating the authors’ recommendations for parameters (providing biological/technical replicates; labeled as TALON_reco in Figure 2) significantly improved precision, which nonetheless falls short of most other tools despite decent sensitivity. Among other reference-guided tools, FLAMES displays poor sensitivity but the assemblies, albeit incomplete, remain accurate, presumably due to a stringent filtering step in the pipeline (see Methods for further details). FLAIR displays reasonable precision but ranks relatively poorly overall in *guided* mode.

**Figure 2.**
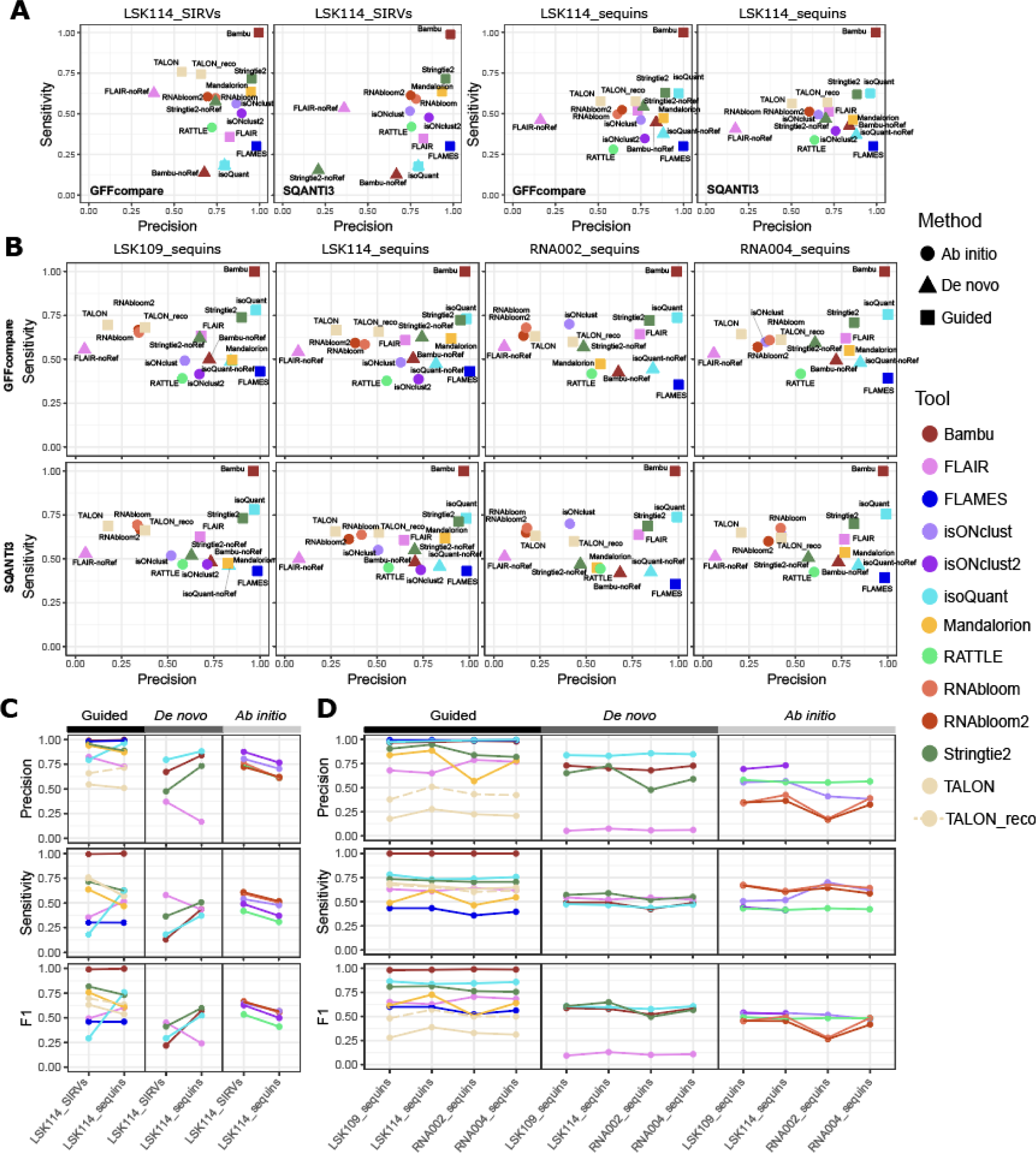
Precision, sensitivity and F1 for all sampled datasets and algorithms. **(A)** Precision and sensitivity values calculated as per GFFcompare and SQANTI3 classifications for assembly outputs from both 40,000 reads from SIRV (left panels) and Sequins™ (right panels) with cDNA+PCR ligation protocol LSK114. **(B)** Precision and sensitivity values calculated as per GFFcompare and SQANTI3 classifications for all Sequins™ dataset assembly outputs available generated from 150,000 input reads. **(C)** Mean precision, sensitivity and F1 values calculated from GFFcompare and SQANTI3 classifications for the same datasets as (A). **(D)** Mean precision, sensitivity and F1 values calculated from GFFcompare and SQANTI3 classifications for the same datasets as (B). For individual values, in (C) & (D) see **Supplemental** Figure 1.

*De novo* methods generally have lower precision and sensitivity compared to *guided* methodologies, even when employing the same software. Here, IsoQuant’s *de novo* approach (labeled as IsoQuant-noRef in **Figure 2**) displays better precision than Stringtie2 but lower sensitivity. Stringtie2 (Stringtie2-noRef) has better overall sensitivity than the other *de novo* tools, especially in cDNA (LSK109 & LSK114 in **Figure 2B**). Although Bambu’s *de novo* mode has a drastic drop in precision and sensitivity when compared to *guided* mode, it performs slightly better than Stringtie2-noRef (especially in dRNA; **Figure 2B**). FLAIR in *de novo* mode regrettably does not instill confidence in any tested modality, highlighting an issue with a very high false positive rate. Interestingly, all *de novo* methods performed relatively poorly compared to *guided* and *ab initio* assembly modalities on the LSK114 SIRV dataset, suggesting that *guided* and *ab initio* strategies might be most suitable for resolving isoforms in genes presenting complex alternative splicing.

The *ab initio* tools RNAbloom and RNAbloom2 present low precision but relatively high sensitivity, akin to the *guided* version of TALON. A sharp eye will notice performance values differ between **Figure 2A** & **2B**, indicating that depth of sequencing can affect assembly precision (see below). The poorest performer among *ab initio* methods appears to be RATTLE, with low sensitivity and precision slightly above 50%. Despite precision values of ∼50-60%, RATTLE outperforms other *ab initio* methods for precision on dRNA, while isONclust2 is the most precise for cDNA (N.B. it does not process dRNA)(**Figure 2B**). Despite the high precision of isONclust2, it has a low sensitivity of ∼50%, while the previous version (isONclust) has a higher sensitivity ranging between 50-75% but lower precision.

Noting the differences between the latest (LSK114) and earlier (LSK109) ONT cDNA library technologies (**Figure 2D**) one can observe the improvement in average precision of the TALON_reco, Stringtie2 (both, the guided and *de novo* versions), RNAbloom and RNAbloom2 with the latest technology, which presents less sequencing errors. The precision and sensitivity shown in **Figure 2C** & **2D** are the mean of values generated from GFFcompare and SQANTI3’s TP, FP and FN classifications, which are shown distinctly in **Supplemental Figure 1**. Similar precision improvements are observed for RNAbloom and RNAbloom2 with the latest version of the dRNA chemistry (RNA004) versus the older version (RNA002). Certain tools, such as Mandalorian and Stringtie2-noRef, have better average sensitivity and precision, respectively, with RNA004 compared to RNA002 (**Figure 2D**). FLAMES is also affected in average sensitivity measures for the RNA002 dataset, although to a lesser extent than the gap between Mandalorian’s performance between the chemistries.

Besides the performance of individual tools, an interesting trend is observed concerning the choice of reference material: Methods applied to SIRV data present a broader range of sensitivity versus a broader range of specificity for the Sequins™ data (**Figure 1A**). This may reflect the different composition of the mixtures, with SIRVs containing relatively less genes (and more isoforms per gene) than Sequins™. Control dependent differences are also observed for individual tools (**Figure 2A,C**). There is a clear decrease in sensitivity and overall F1 score for isoQuant (*guided*) when applied to the SIRV data, with Flair (*guided*) following a similar pattern. Mandalorion, Stringtie2, TALON and TALON_reco seem to have improved sensitivity and F1 score with SIRV versus Sequins™. Among the *de novo* strategies, we can observe a dramatic decrease in the precision and sensitivity of Bambu, isoQuant and Stringtie2 for the SIRV dataset, while Flair follows the opposite trend (corroborated in **Figure 2A,C**). There is an improvement in performance of all *ab initio* tools when used on the SIRV dataset (**Figure 2C**; absolute values in **Supplemental File 1**), suggesting that different transcriptomic features will globally influence accuracy. This is exemplified with RNAbloom and RNAbloom2, whose accuracies are highly influenced by input parameters. N.B. TALON and TALON_reco pipelines produced no results when evaluated on SIRV with SQANTI3 (**Figure 1A**) due to issues with SQANTI3 handling of the strandedness in the assembly.

Overall, the F1 score of reference-guided tools remains fairly consistent across datasets except IsoQuant, which performs a lot better on Sequins™ than SIRVs (75% and 25%, respectively; **Figure 2C**). We observed that *de novo* and *ab initio* methods have a very similar predictive performance overall (average F1 score, **Figure 2C,D**), with RNA002 and FLAIR as notable exceptions.

Significant differences in performance were observed between many assembly tools that can be attributed to the method used for qualitative evaluation. Stringtie2-noRef performs at 55% and 75% in sensitivity and precision, respectively, when evaluated with GFFcompare but both metrics fall to 15% with SQANTI3 for the same assembly (**Supplemental Figure 1A,B**). This suggests that the choice of quality metrics used to benchmark assembly and isoform detection performance can have a drastic impact on the resulting assembly strategy, while highlighting that isoform class definitions is a somewhat subjective evaluation.

These comparisons were performed with static read depths to directly compare assembly accuracies in a normalized manner. Since depth of sequencing can impact assembly accuracy by increased sampling of lower abundance transcripts and experimental artifacts, we evaluate the performance of representative tools on different sampling depths of the LSK114_sequins dataset, from 1,000 reads to 4.3 million reads (**Figure 3**). Overall, higher depth of sequencing is associated with increased sensitivity and lower specificity, with the notable exception of *guided* assembly modalities. Bambu, in particular, outputs all transcripts from the reference, yielding a sensitivity above the maximal theoretical value. Both IsoQuant and Stringtie2 approach 100% sensitivity and 75% precision at maximal depth, irregardless of the scoring metric. However, the *de novo* versions of Bambu and IsoQuant stagnate after 100k and 500k reads, respectively, evolving within the 48-55% sensitivity and 60-75% precision ranges (**Figure 3**, second column). In contrast, *de novo* Stringtie2’s sensitivity scales with sampling depth. Of the two *ab initio* methods we tested, isONclust2 presents relatively stable precision in function of depth, with comparable if not superior performance to the 3 *de novo* methods in the middle panel. A drastic increase in precision for the isONclust result at 500k reads, evaluated using SQANTI3, presents an unusual outlier that we could not dismiss. Overall, the surveyed *ab initio* and *de novo assembly* modalities display a significantly lower sensitivity than theoretically expected, suggesting that the use of a minimum coverage threshold might restrict isoform discovery. Interestingly, *ab initio* methods appear to perform slightly better than *de novo* strategies when evaluating with SQANTI3 (at least, for sensitivity), while the converse is observed with GFFcompare.

**Figure 3.**
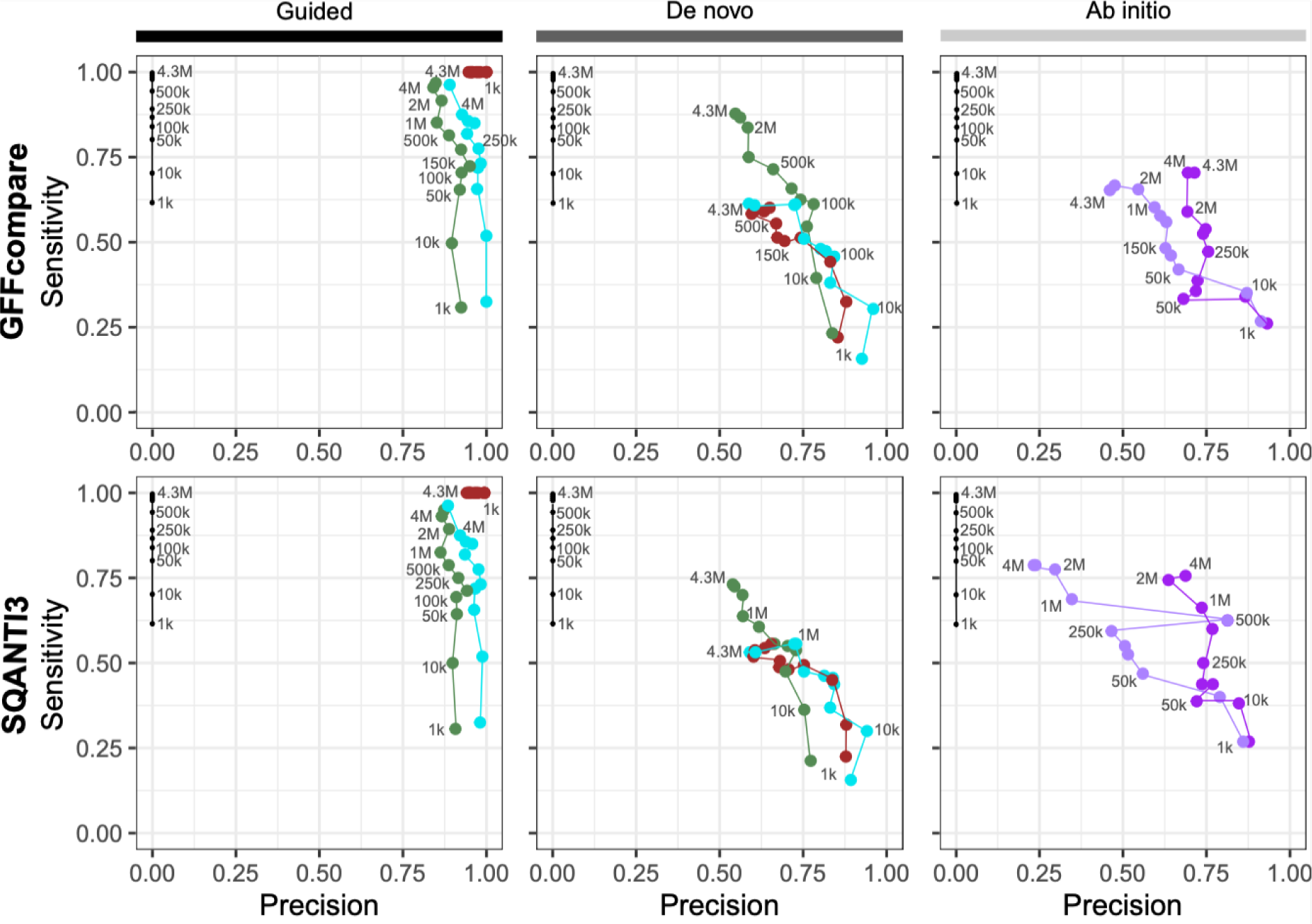
Sequencing depth impact on assembly. Precision and sensitivity values calculated as per GFFcompare (top panels) and SQANTI3 (bottom panels) classifications for all subsets available (ranging from 1,000=1k to 4,300,000=4.3M reads) of the LSK114_sequins dataset. Selected pipelines are Bambu, isoQuant and Stringtie2 for guided strategies; Bambu-noRef, isoQuant-noRef and Stringtie2-noRef for *de novo* strategies; and isONclust and isONclust2 for *ab initio* assembly strategies. The “max value per relative abundance” is calculated, from expected abundance as disclosed by Sequins™, as the maximum sensitivity for the number of transcripts detectable with at least 1 transcript per million (TPM) based on the number of input reads.

One of the advantages of generating bespoke transcriptome annotations is to improve isoform-level quantification, thereby providing a robust foundation upon which subsequent post-processing procedures can be reliably executed. We compared the expected vs observed quantifications obtained based on the assemblies generated by Bambu, IsoQuant, Stringtie2-noRef and Flair-noRef. These tools have been selected to highlight biases in assembly precision or sensitivity and their potential effect on resulting quantification. Bambu and IsoQuant have a similar precision (99% and 98% respectively) but a gap of 26% in sensitivity, whilst Stringtie2-noRef and Flair-noRef have a similar sensitivities (62% and 55% respectively) but a gap of 65% in precision (**Figure 2B-D**). We only assessed correlations for assembled transcripts that were classified as true positives while using the complete assembly as a reference for quantification (see Methods). Results indicate that *guided* strategies (i.e. Bambu and IsoQuant) are very close to the expected abundances (R^2^ > 0.8) and stable across evaluators (**Figure 4**). However, although Stringtie2-noRef and FLAIR-noRef yield similar correlation coefficients (between 0.69-0.76 and 0.6-0.76 respectively), the FLAIR-noRef quantification generates a regression curve substantially lower than the expected x=y line, indicating that erroneous transcripts might be ‘absorbing’ read assignments from true positive isoforms. As expected given the lack of reverse transcription and PCR, direct RNA reads (RNA002 & RNA004) generate quantifications that correlate better with expected values than cDNA for all assemblies.

**Figure 4.**
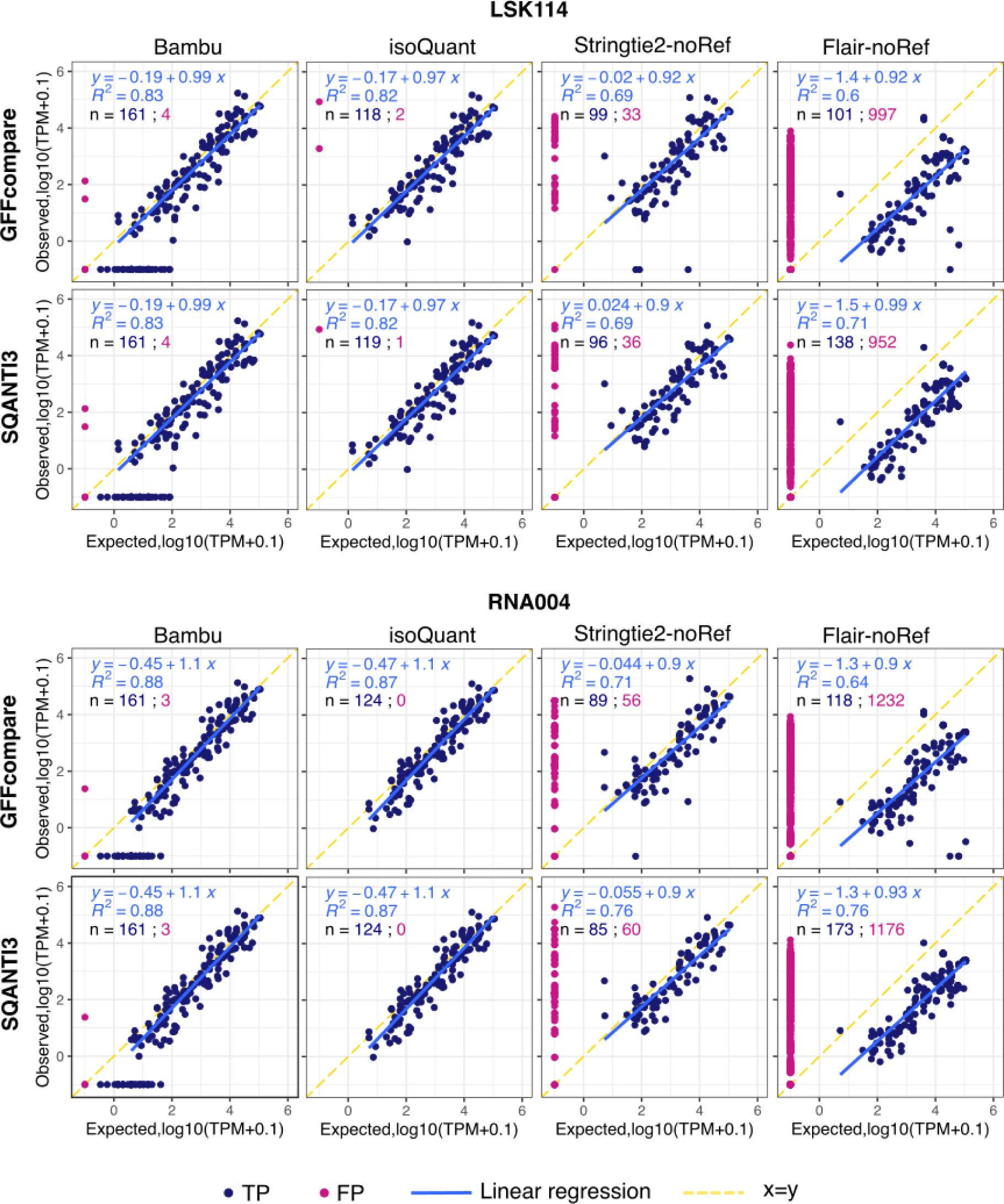
Assembly impact on quantification. Transcript-level quantification results as generated by Salmon from read alignments to transcriptome assemblies generated by Bambu, isoQuant, Stringtie2-noRef and Flair-noRef pipelines outputs on the LSK114_sequins_sub150k and the RNA004_sequins_sub150k datasets. Only True Positive (TP, see Methods for definition) transcripts as defined for GFFcompare (top panels) and SQANTI3 (bottom panels) are assigned to an expected abundance, as per abundance information provided by Sequins™. False Positive (FP) transcripts were assigned a null expected abundance and removed from the linear regression equation and correlation coefficient (R^2^) calculation. Number of transcripts for each category (TP, FP) is available on each panel as n = [number of TP transcripts in assembly] - [nb of FP transcripts in assembly], as well as the correlation coefficient.

With respect to ease of use and the execution time, most (but not all) tools leverage CPU multithreading (20 threads applied when possible). Depending on the input file, some *guided* methods (FLAIR, FLAMES, IsoQuant, Mandalorion) integrate the alignment step, while others do not (TALON, Stringtie2, Bambu). As shown in **Figure 5**, TALON_reco displays a significantly shorter alignment step and a significantly longer pre-processing step due to the alignment options and the use of the additional TranscriptClean script, which consumes 98% of the pre-processing time. Otherwise, all assemblers require similar computation time for 150 k reads except FLAIR, which, like TALON, includes a correction step. Notably, FLAMES performs similarly to other *guided* methods despite using a single thread. For *de novo* and *ab initio* methods, IsoQuant-noRef (13min15sec), isONclust (7min) and isONclust2 (1min30sec) stand out with significantly shorter execution times than all the other tools.

**Figure 5.**
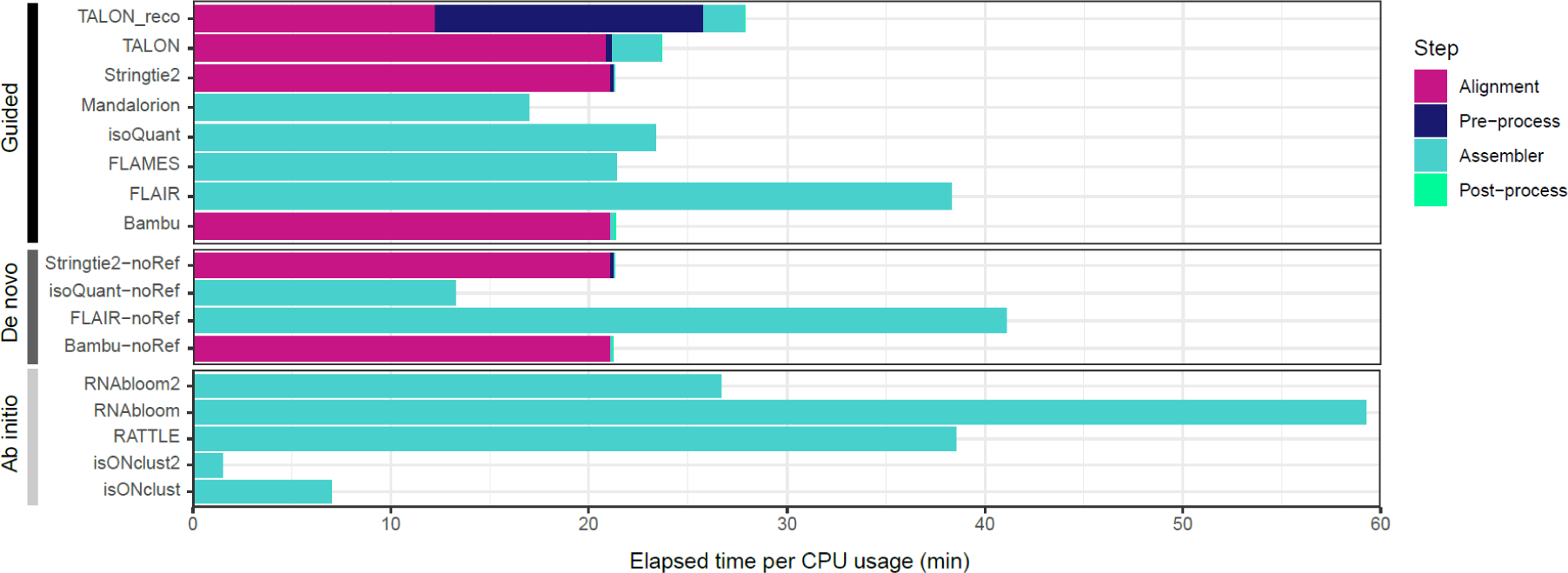
Execution time. Elapsed time as outputted by the /user/bin/time system function multiplied by the CPU usage percentage for all steps of all pipelines applied to the LSK114_sequins_sub150k dataset. When allowed, all tools were executed with 20 threads on a workstation with 2x16 core 2.30 GHz Intel Xeon Gold 5218 workstation with M.2 NVMe solid state drives.

## Discussion

This benchmarking study on assembly algorithms using nanopore RNA-seq data was originally intended to expose the best methodology for the resolution of RNA isoforms. This was prompted by improved nanopore sequencing accessibility and accuracy that can easily identify numerous novel RNA isoforms, even in extensively sequenced model organisms. In our opinion, the common practice of relying on a reference transcriptome annotation (i.e., Refseq or Gencode) can lead to significant biases by including genes and isoforms that are not expressed in a given experimental condition and, more importantly, by ignoring or omitting unannotated transcripts that might be highly expressed or significantly regulated between conditions. This is notably the case for extensively spliced long non-coding RNA genes, which now outnumber protein-coding genes in recent Gencode annotations. The results we have obtained from RNA standards indicate that there is substantial variability in the accuracy of isoform reconstruction tools; a greater diversity than we expected given the single molecule nature of nanopore sequencing data. Hence, there appears to be no clear “one size fits all” solution for transcriptome assembly at the moment.

There are nonetheless several notable observations that can help guide future nanopore RNA-seq analyses. Providing a reference annotation and a reference genome unsurprisingly yields the most accurate (although not necessarily perfect) results. This modality might not detect novel isoforms beyond those in the reference–a possibility we briefly explored by removing a fraction of the isoforms in the reference annotation and assessing their discovery thereafter (not shown). In practice, this equates to the *de novo* modality, where reads are aligned to a genome prior to assembly and without a reference. Although this strategy greatly facilitates isoform discovery, the assembly must then be annotated by comparison to a gene reference with tools like GFFcompare and SQANTI3, which is a somewhat subjective undertaking that can complexify downstream analyses (see below). For *de novo* assembly, IsoQuant consistently demonstrates the least amount of false positive transcripts across all surveyed datasets, while StringTie2 might provide the greatest overall accuracy (F1 score), closely followed by reference-free Bambu and IsoQuant (**Figure 2**). Surprisingly to us, the performance of *ab inito* methods compares favorably to *de novo* techniques in accuracy and execution time, especially isONclust & isONclust2. However, isONclust2 requires some enhancements to accommodate dRNA reads and to facilitate its integration into a streamlined analytical workflow. These *ab initio* techniques are often overlooked in model organisms with high-quality genomes, as well as many benchmarking studies. Their assembled transcriptome output is typically aligned to a genome prior to comparative reference gene annotation, thus also requiring tools such as GFFcompare and SQANTI3. Our analysis concurs with the conclusions of recent studies on the accuracy, effectiveness and versatility of Bambu, IsoQuant and Stringtie2 (Dong et al. 2023; Pardo-Palacios et al. 2023b). However, our assessment contradicts conclusions by the LRGASP consortium on the utility of FLAIR, found to be one of the most robust strategies for both well annotated organisms and orthogonal data integration. Our results suggest that the accuracy of the versions we tested lies far behind other tools, particularly when executed without a reference annotation (**Figure 2**).

Given the unsurprising tradeoff between sensitivity and specificity, some methods should be privileged over others depending on the objective of the sequencing experiment. For example, when seeking to identify new isoforms of known genes, certain tools might not be suitable. This is the case for IsoQuant and Bambu in *de novo* mode on SIRVs (sensitivity and specificity of ∼15%, **Figure 2A**), a spike-in control set with fewer genes and more alternative splicing isoforms than Sequins™, where these tools performed much better (45-55% sensitivity). It should be mentioned that the depth of sequencing may also play a role in assessing each method’s perceived accuracy, as we demonstrated in **Figure 3**. Unfortunately, the limited SIRV and dRNA sequencing depth and the large number of surveyed tools restricted a more comprehensive evaluation of the impact of depth on assembly accuracy, which undoubtedly has an impact on the assessment of true and false positive isoforms.

Certain factors that can affect transcriptome assembly were not directly evaluated in this study, either by design or practicality. Firstly, base calling algorithms and their associated error rates could potentially impact alignment and assembly accuracy, notably at exon-intron boundaries. We applied the latest base calling algorithms when preparing this manuscript; older versions (with high error rates) might produce different results. Secondly, we employed a single pairwise alignment tool (minimap2) when alignment was required. Alternative aligners could significantly impact assemblies, particularly in *guided* and *de novo* modalities. The impact of alignment can be major––when confronted with fragmented reads, alignment algorithms can force terminal sequences spanning an exon junction to align within the intron as the affine gap penalty may be substantially larger than the associated mismatch penalty, leading to an apparently longer ‘read-through’ exon, a common observation in de novo transcriptome assemblies. Thirdly, the choice of reverse transcription enzymes, elongation times and PCR cycles can also have a significant impact on the results; the manufacturer’s recommended protocols at the time of data generation were used. An additional experimental variable could be freeze-thaw cycles of the spike-in controls. To the best of our knowledge, only fresh controls were used. Fourthly, we used the default or recommended execution parameters for the assembly algorithms. It is to be expected that tweaking these would significantly impact results. In particular, we did not attempt to normalize the minimum coverage threshold when this parameter was available, defaulting to the developers’ recommended settings. It is to be expected that a lower threshold would increase sensitivity and, potentially, reduce specificity for a given tool. In general, the developers bear the responsibility of optimal default parameter settings. Lastly, we did not evaluate the relative impact of transcript and sequence features on assembly accuracy. GC content, transcript length, exon length, number of exons, isoform diversity per gene are features that might help pinpoint the strengths and limitations of certain tools, while guiding algorithmic improvements to distinguish between biological signal and noise.

This study substantiates that assembling a long read transcriptome–even from relatively simple, well-defined spike-in controls–remains a complex and error-prone endeavor. Even the methods to compare and evaluate transcriptome annotations are subject to variation, exposing the subjective nature of qualitative transcript classifications. Most studies rely on either GFFcompare (Stuart et al. 2024; Saha et al. 2024) or SQANTI3 (Pardo-Palacios et al. 2023b; Brooks et al. 2024) to this avail. When comparing both methods on the same data, we observed surprisingly different assembly accuracies (c.f. the performance of Stringtie2-noRef with SIRV data in **Figure 2**, **Supplemental Figure 1**). While GFFcompare assesses a *de novo* transcriptome assembly with 75% precision and 55% sensitivity; SQANTI3 assesses 15% for both. This can partially be explained by what transcript classifications we esteemed to define true or false positives (we considered incomplete transcripts, ‘c’ for GFFcompare and ‘ISM’ for SQANTI3, as true positives) but also by the way both tools classify transcripts, such as nuances in the assessment of intron chains or splice junction mappings. Another likely source of divergence is how GFFcompare systematically selects the most similar reference transcript to qualify an isoform, while SQANTI3 remains impartial and simply classifies any isoform that doesn’t perfectly match the reference features as “new”. Nonetheless, the isoform definitions we employed herein form an arguably more stringent approach compared to other studies, where all transcripts classifications that overlap the reference genes are considered true positives (Song et al. 2019). An interesting solution lies in the sequential use of GFFcompare and SQANTI3 annotation evaluators described by Wijeratne et al. (Wijeratne et al. 2024). The authors use GFFcompare to compare transcriptome features across samples and generate a combined list of non-redundant isoforms, while SQANTI3 is employed for comprehensive characterization of long-read transcript sequences, including quality control, identification, and quantification of full-length transcripts.

In summary, evaluating the accuracy of long-read transcriptome assembly strategies exposes the multi-faceted considerations that impact isoform-level analyses. In addition to the choice of assembly modality and algorithms, the nature of queried sequencing data and assembly evaluator characteristics impose nuanced trade-offs for isoform discovery and transcriptome annotation. The most important consideration, however, is the inclusion of well defined spike-in controls to help quantify artifacts of transcriptome assembly and establish a baseline of truthfulness in the output. By integrating these insights, researchers can make informed choices in selecting assembly methods tailored to their specific experimental objectives and resource constraints.

## Methods

### Samples sequencing and data acquisition

The following datasets, LSK109_sequins and LSK114_sequins were obtained by preparing 15ng of sequin standards RNA mix A (Hardwick et al. 2016) following the PCS109 library preparation protocol as recommended by manufacturer [Oxford Nanopore Technologies©, UK] up until the PCR step. Then, the protocol LSK109 (amplicons) and LSK114 (amplicons) was applied to the resulting cDNA. 50 fmol and 20 fmol of the respective final libraries were loaded onto a R9.4.1 flowcell (flowcell ID: FAN54376) and a R10.4.1 flowcell (flowcell ID: FAV99142) and ran for 72 hours on a GridION sequencer and 12 hours on a PromethION sequencer, respectively.

The LSK114_SIRVs dataset was obtained by preparing 0,0375ng of SIRV set-04 [Lexogen, Austria] spiked-in with 125ng of a human RNA sample, as recommended by the manufacturer. As for previous datasets, samples were processed following the PCS109 library preparation protocol up until the PCR step; then the protocol LSK114 (amplicons) was applied to the resulting cDNA as recommended by the manufacturer [Oxford Nanopore Technologies©, UK]. 18 fmol of the final library was loaded onto a R10.4.1 flowcell (flowcell ID: PAK95982) that ran for 72 hours on a PromethION sequencer. A reload was done with an additional 18 fmol of the library after 29 hours. Only reads mapping to the SIRV set-04 genome were retrieved using minimap2 and seqtk toolkit (https://github.com/lh3/seqtk).

For the RNA002_sequins dataset, 300ng of sequin standards RNA mix A (Hardwick et al. 2016) was prepared following the RNA002 library preparation protocol as recommended by the manufacturer [Oxford Nanopore Technologies©, UK]. 20ng of the final library was loaded onto a R9.4.1 flowcell (flowcell ID: PAM58324) and ran on a PromethION sequencer for 72 hours.

RNA004_sequins dataset was acquired by preparing 4 samples with 1.5μg of human spiked-in with 15ng of sequin standards RNA mix A (Hardwick et al. 2016). Samples were prepared following the RNA004 protocol (bêta testing phase: V2 April 2023). 20ng of the final libraries for each sample was loaded onto individual RP4_bêta-testing flowcells (flowcell IDs: PAO83093 - PAO83456 - PAO84072 - PAO96683) and ran for 68 hours on a PromethION sequencer. A reload with the same amount of library was done after 44 hours and 15 minutes of sequencing. Only reads mapping to the Sequins™ reference genome were retrieved using minimap2 and seqtk toolkit (https://github.com/lh3/seqtk).

### Datasets pre-processing

All datasets were basecalled with Guppy v6 [6.0.1 to 6.5.7] super high accuracy mode. For cDNA datasets (LSK109_sequins, LSK114_seuins, LSK114_SIRVs), pass and fail reads from basecalling were trimmed and reoriented using Pychopper v2.7.9 (https://github.com/epi2me-labs/pychopper). Full-length and rescued reads from Pychopper were then processed like raw basecalled pass/fail reads for dRNA datasets (RNA002_sequins, RNA004_sequins). Minimap2 v2.24 (Li 2021) was used to generate .bam, and .sam files (-ax splice --secondary=no). The option --MD and .sam output was used for TALON pipeline only, as required by the tool.

### Subsampling and transcriptome assembly

The five datasets (PCS109_sequins, LSK114_sequins, LSK114_SIRVs, RNA002_sequins, RNA004_sequins) were randomly subsampled after pychopper pre-porcessing for cDNA datasets and after base calling for dRNA datasets to have the following amount of reads: 1k - 10k - 50k - 100k - 150k - 250k - 500k - 1M - 2M - 4M - Total. All subsampled datasets were processed through assembly as described below. Subsampled datasets are referred to as subX, where X is the number of reads in the dataset.

Each tool was run with default parameters using 20 threads when possible. Some tools were executed in two different ways (Stringtie2/Stringtie2-noRef, Bambu/Bambu-noRef, Flair/Flair-noRef, isoQuant/isoQuant-noRef) when assembly with and without a transcriptomic reference was allowed. For RNAbloom, the final assembly assessed doesn’t contain the assembled short transcripts. isONclust2 was applied to cDNA datasets only (as it is not yet available for dRNA). FLAMES .gff3 output is filtered based on its transcripts evaluation provided : only transcripts classified as “true” in the ‘isoform_FSM_annotation.csv’ file are filtered and considered as final assembly. Further details of pipelines and their execution is available on github (https://github.com/msagniez/LRassBench).

**Table 2.**
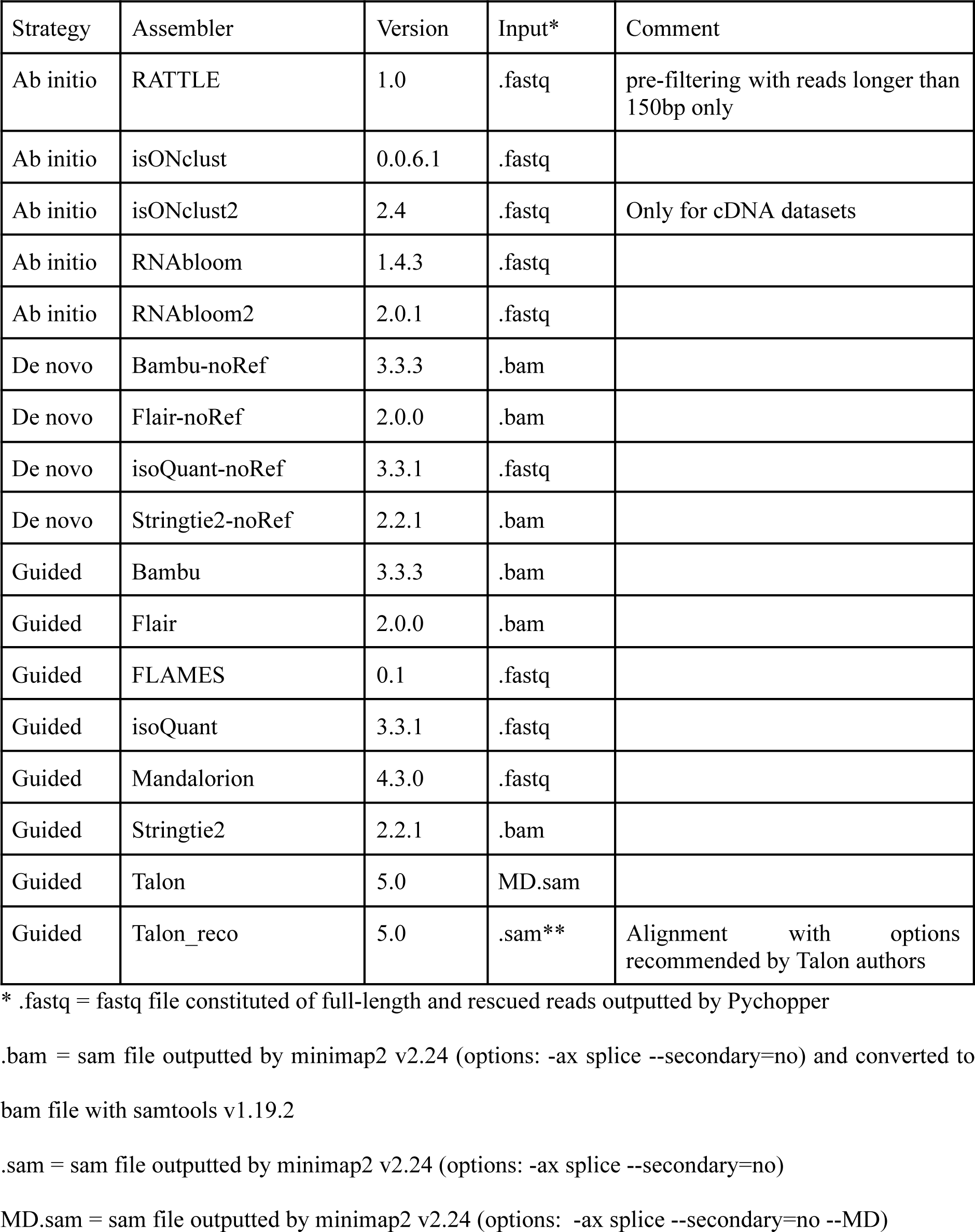
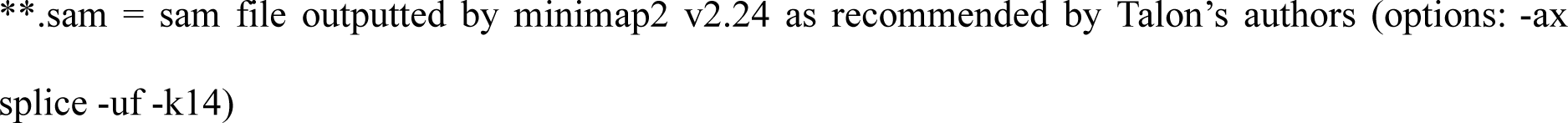
List of tools and corresponding input files.

| Strategy | Assembler | Version | Input* | Comment |
| --- | --- | --- | --- | --- |
| Ab initio | RATTLE | 1.0 | .fastq | pre-filtering with reads longer than 150bp only |
| Ab initio | isONclust | 0.0.6.1 | .fastq |  |
| Ab initio | isONclust2 | 2.4 | .fastq | Only for cDNA datasets |
| Ab initio | RNAbloom | 1.4.3 | .fastq |  |
| Ab initio | RNAbloom2 | 2.0.1 | .fastq |  |
| De novo | Bambu-noRef | 3.3.3 | .bam |  |
| De novo | Flair-noRef | 2.0.0 | .bam |  |
| De novo | isoQuant-noRef | 3.3.1 | .fastq |  |
| De novo | Stringtie2-noRef | 2.2.1 | .bam |  |
| Guided | Bambu | 3.3.3 | .bam |  |
| Guided | Flair | 2.0.0 | .bam |  |
| Guided | FLAMES | 0.1 | .fastq |  |
| Guided | isoQuant | 3.3.1 | .fastq |  |
| Guided | Mandalorion | 4.3.0 | .fastq |  |
| Guided | Stringtie2 | 2.2.1 | .bam |  |
| Guided | Talon | 5.0 | MD.sam |  |
| Guided | Talon_reco | 5.0 | .sam** | Alignment with options recommended by Talon authors |

\* .fastq = fastq file constituted of full-length and rescued reads outputted by Pychopper
.bam = sam file outputted by minimap2 v2.24 (options: -ax splice --secondary=no) and converted to bam file with samtools v1.19.2
.sam = sam file outputted by minimap2 v2.24 (options: -ax splice --secondary=no)
MD.sam = sam file outputted by minimap2 v2.24 (options: -ax splice --secondary=no --MD)
\*\*sam = sam file outputted by minimap2 v2.24 as recommended by Talon's authors (options: -ax splice -uf -k14)

### Assembly quality assessment

Final assemblies were mapped to the original reference transcriptomes using two different tools to define the mappings: GFFcompare (Pertea and Pertea 2020) (.tmap file) and SQANTI3 (Pardo-Palacios et al. 2023a) (_classification.txt file). All Figures were plotted using the ggplot2 library in R.

### Sensitivity, Precision and F1 calculation

As described in Figure 1, true positive transcripts (TP) are transcripts labeled “=” and “c” for GFFcompare as well as “Full Splice Match” and “Incomplete Splice Match” (except intronic transcripts) for SQANTI3. Potential new transcripts are the transcripts labeled “u” by GFFcompare and “Intergenic” by SQANTI3. These potential new transcripts are excluded from true positive, false positive and false negative sets. False positive transcripts (FP) are the ones that don’t match the labels cited above. False negatives (FN) are the transcripts found in the reference but not assembled by the tools. The sensitivity is calculated as *TP*/(*TP* + *FN*) and the precision is calculated as *TP*/(*TP* + *FP*). The F1 score is calculated as *F*1 = (2 * *Precision* * *Sensitivity*)/(*Precision* + *Sensitivity*).

### Transcripts discovery accuracy assessment

We selected 9 transcripts to remove from the original Sequin™ .gtf annotation (R1_13_1 ; R1_21_2 ; R1_51_1 ; R2_116_1 ; R2_116_2 ; R2_47_2 ; R2_59_3 ; R2_6_2 ; R2_72_1) based on their theoretical abundance, length and number of isoforms. All guided tools were executed again using LSK114_sequins_sub150k dataset and the incomplete .gtf annotation. The GTFs from all the tools were parsed in R, and all the genomic coordinates of the assembled transcripts were plotted. The data was used to narrow down the assemblies that mapped to the region of the 9 artificially created ‘novel’ transcripts. This subset was then filtered for exons that completely mapped to the known exons, as we wished to retain the ones that mapped to the nine novel isoforms only. This filtered dataset was used to visualize the assemblies, compared to the nine novel isoforms, using ggplot2 and ggtranscript (Supplemental figure 1 and 2).

### Sequencing depth analysis

We show Precision and Sensitivity results for the 5 most performant tools (Bambu, isoQuant, Stringie2, isONclust, isONclust2) on all subsets of the LSK114_sequins dataset. For each of these subsets we also calculated the maximum sensitivity per relative abundance (MaxSensitivity) considering the theoretical abundance of each transcript in the mix as:

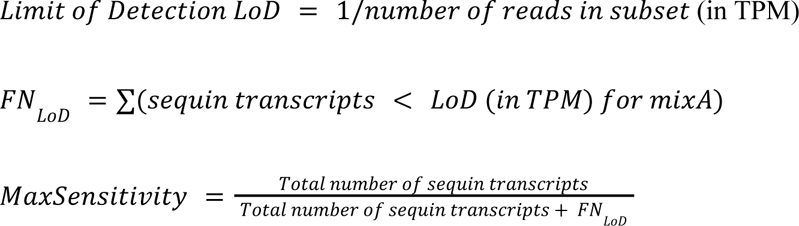

### Quantification

Quantification was obtained after mapping the original .fastq files of datasets LSK114_sequins_sub150k and RNA004_sequins_sub150k to the corresponding output assembly of Bambu, isoQuant, Stringtie2-noRef and Flair-noRef with minimap2 v2.24 [options: -a -N 100] and samtools v1.19.2 to convert to bam. Salmon v1.10.1 (Patro et al. 2017) was then executed. Experimental quantifications (TPM) of TP (as described in Sensitivity, Precision and F1 calculation section) were compared to theoretical abundances converted to TPMs to generate Pearson’s correlation coefficient. TPs with null experimental quantifications were excluded from the coefficient calculation, as they are considered not found in assembly. FP transcripts were attributed a theoretical abundance of 0, as these are not real transcripts and should not be present.

## Data access

Assembly annotations evaluated as well as GFFcompare (.tmap file) and SQANTI3 (_classification.txt file) evaluations are available along with the code used in our github page : https://github.com/msagniez/LRassBench. Input .fastq files as used in this study for SIRV and sequin data are available on SRA [accession numbers]

## Acknowledgments

We would like to thank Eduardo Eyras and Richard Kuo for comments and feedback during the preparation of this work; David Barda and Tim Mercer from SequinsSequin™ who donated spike-in controls essential for this work; Libby Snell and Etienne Rainmondeau from Oxford Nanopore Technologies for support with RNA004 direct RNA beta testing; and the developers and bioinformatics community that contributed to the various open-source algorithms that were tested.

This work was supported by a Fonds de recherche du Québec–Santé Junior 1 fellowship [project 284217], a Canadian National Science and Engineering Research Consortium Discovery grant [DGECR-2022-00207], a Cole Foundation Transition Award, a Canadian New Frontiers in Research Fund Exploration Grant [NFRFE-2021-01005], by the Canadian Foundation for Innovation John R. Evans Leaders Fund [project 40767] to MAS, and by a Cole Foundation PhD fellowship to MS. Computing resources were partially provided by a research allocation from the Digital Research Alliance of Canada and Calcul Québec.

## Author contributions

MAS conceived the study and secured funding. MS, BP, MR and MAS generated sequencing libraries and generated sequencing data. MS, AB, SMS & CVO performed bioinformatics analyses. MS & AB generated Figures. MS, AB & MAS wrote the manuscript with input from all authors.

